# Global spatial potential for implementing land-based climate mitigation

**DOI:** 10.1101/2024.01.04.574063

**Authors:** Evelyn M. Beaury, Jeffrey Smith, Jonathan M. Levine

## Abstract

Land-based mitigation strategies (LBMS) are critical to reducing climate change and will require large areas for their implementation. Yet few studies have considered how and where LBMS compete for land or are mutually compatible across Earth’s surface. We derived high resolution estimates of the spatial distribution of 19 different LBMS. We estimated 8.9 billion ha suitable for LBMS across the Earth, including 5.6 Bha suitable for only one of the studied strategies and 3.3 Bha suitable for multiple LBMS. We identified significant conflicts between better carbon management of existing land cover types, and restoration-based strategies such as reforestation. At the same time, several agricultural management LBMS were compatible over large areas, including for example, enhanced weathering and improved plantation rotations. Our analysis presents local stakeholders, communities, and governments with the range of LBMS options, and the opportunity costs associated with scaling up any given LBMS to reduce global climate change.

---

Limiting climate warming to 2°C requires a dramatic energy system transformation paired with the widespread deployment of land-based mitigation strategies (LBMS) – a suite of land use and land management practices that harness plant and soil processes to reduce carbon emissions and increase carbon sequestration^1–5^ (Table 1). While cutting carbon emissions is the single most important component of reaching net zero, these cuts must be complemented by LBMS as they, unlike emissions reductions alone, allow us to remove emitted CO_2_ from the atmosphere^1,6^. To meet their full potential, however, LBMS must be implemented across large spatial extents^6–8^, requiring major shifts in Earth’s distribution of land cover. The degree to which LBMS can address climate change therefore depends on the extent of land available for, and in turn dedicated to implementing land-based climate mitigation. Accordingly, identifying the spatial distribution of opportunities for LBMS is particularly critical for actualizing national and global commitments to reduce climate change^9,10^.

**Table 1.**
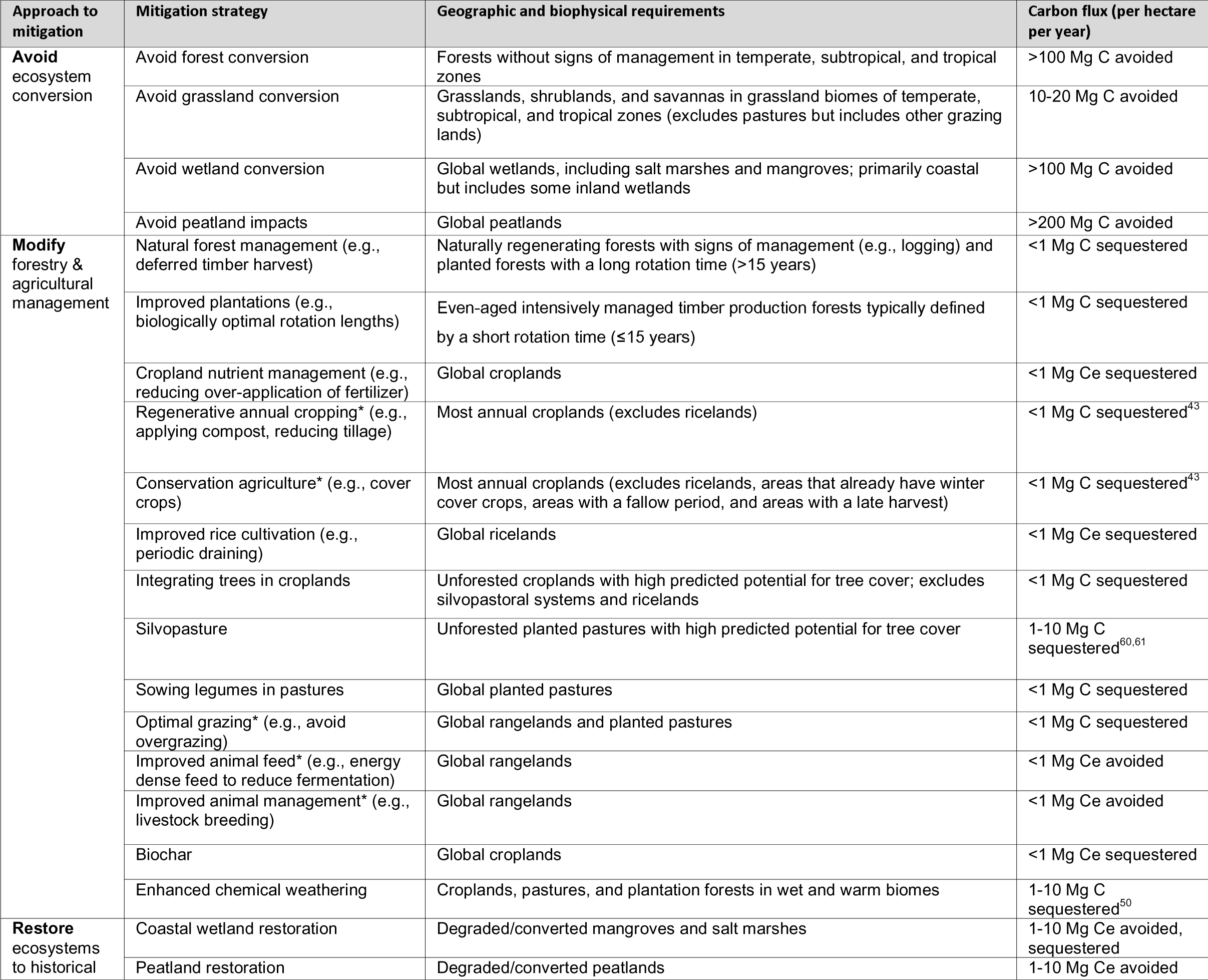
Land-based climate mitigation strategies, a brief description of their geographic requirements, and a rounded estimate of the intensity of carbon emissions avoided or additional carbon sequestered per hectare per year. The latter is intended as a means of comparison across the strategies, but large uncertainties remain in the magnitude, temporal, and spatial variation in these estimates. Carbon flux was adapted from Griscom et al. (2017), unless otherwise noted. 19 of the strategies listed here were mapped in our analysis (all but those with an asterix). Additional details are provided in SI.

In recent years, approaches to land-based mitigation have expanded to include a diversity of strategies that differ greatly in their impacts on atmospheric carbon, biodiversity, and human livelihoods^1–3,11–16^. While the growing number of LBMS provides multiple options for addressing global climate change, uncertainties about the efficacy of any one solution vary substantially^17–20^, and little is known about how scaling up the deployment of one LBMS might affect the potential of others. For example, most previous work has focused on mapping the mitigation potential of a single strategy^21–23^ or estimating the maximum area available to multiple LBMS without considering where this land is distributed^1,3^. Studies on LBMS are often further siloed depending on whether mitigation stems from traditional approaches to better land stewardship (e.g., habitat restoration) or from novel modifications to land processes (e.g., enhanced chemical weathering)^7,24,25^. Regardless of this distinction, all of these strategies require land^3,26,27^ and must be deployed in concert to achieve net zero^28^, and thus an approach that quantifies how all LBMS overlap is ultimately needed.

Such a holistic approach is particularly important to avoid overestimating the combined contribution of different LBMS to climate mitigation. For example, agricultural lands are candidate areas for expanding carbon smart management methods as well as for restoring forest^29,30^. However, since these strategies are incompatible on the same landscape, independently considering land available to each LBMS risks overestimating their respective mitigation potential. Similarly, only by overlaying the land available to each strategy can we properly estimate the opportunity cost associated with deploying one alternative versus the other. Conversely, if mitigation strategies with overlapping land requirements are compatible with one another (e.g., enhanced chemical weathering can supplement bioenergy crop production and other sustainable crop management actions^31^), these areas may provide opportunities for amplifying carbon removals. In either case, considering which and how many LBMS are possible across different landscapes can inform society’s choices when using land to meet climate targets.

Ultimately, most decisions to implement a climate mitigation strategy will be made at a local level based on a variety of considerations beyond carbon storage, such as biodiversity conservation, economic feasibility, food security, and more^5,15^. But if local decisions are made independently, and without the broader spatial context for where LBMS could alternatively be deployed, we may miss an opportunity to strategically scale up climate mitigation. In particular, a better understanding of the spatial overlap of different LBMS is necessary for predicting how policies that incentive one mitigation strategy influence the deployment of others^8,9^. Indeed, coordinating the local deployment of the various strategies is essential to meeting national and global climate targets^32,33^, and thus national and global syntheses are needed to inform optimal outcomes for climate, people, and biodiversity.

To meet this need, we synthesized the literature to identify the extent and overlap of land compatible with 19 different LBMS. These strategies (Table 1) help stabilize the climate by (1) avoiding the loss of carbon-dense ecosystems, (2) restoring carbon-dense ecosystems, (3) modifying agricultural and forestry management practices to reduce emissions and increase sequestration, or (4) converting habitat to store additional carbon in aboveground biomass (e.g., afforestation). For each LBMS, we derived high resolution (∼1km) global maps of the area where each mitigation strategy could be implemented across Earth’s surface. Given uncertainties in the socioeconomic factors that may influence the feasibility and efficacy of mitigation^2,5,17–20^, we map LBMS based on their broad biophysical and geographic requirements, using reasonable, strategy-specific constraints (Table 1, SI). We then compared these spatial distributions to identify the number of LBMS that are possible in any one location, including hotspots of climate mitigation opportunities and potential conflicts in the space-use of LBMS.

Specifically, we asked 1) How much area is available for implementing each climate mitigation strategy given its global geographic and biophysical requirements? 2) How many mitigation strategies overlap with one another and where are these overlaps most pronounced? and 3) Which mitigation strategies most frequently compete for space, and which are mutually compatible with one another?

## Results

The global spatial potential for implementing the 19 LBMS varies substantially (Figure 1), with some of the most efficient mitigation strategies constrained to small geographic areas (e.g., avoided peatland impacts and peatland restoration). Due to their biophysical constraints, LBMS are non-randomly distributed across land cover types, continents, and climate zones (Figure S1-19). For example, enhanced chemical weathering can occur throughout temperate and tropical zones but is concentrated in biomes with high precipitation (Figure 1a, e.g., tropical China and India). For bioenergy with carbon capture and storage (BECCS), potential is highly constrained by the location of sedimentary basins for CO_2_ storage, with low known potential for carbon capture and storage (CCS) in Africa or Australia (Figure 1b). In croplands, integrating tree cover while maintaining food production is environmentally suitable in many regions of the Earth (Figure 1c), with the highest density of potential in the midwestern United States and temperate Europe.

**Figure 1.**
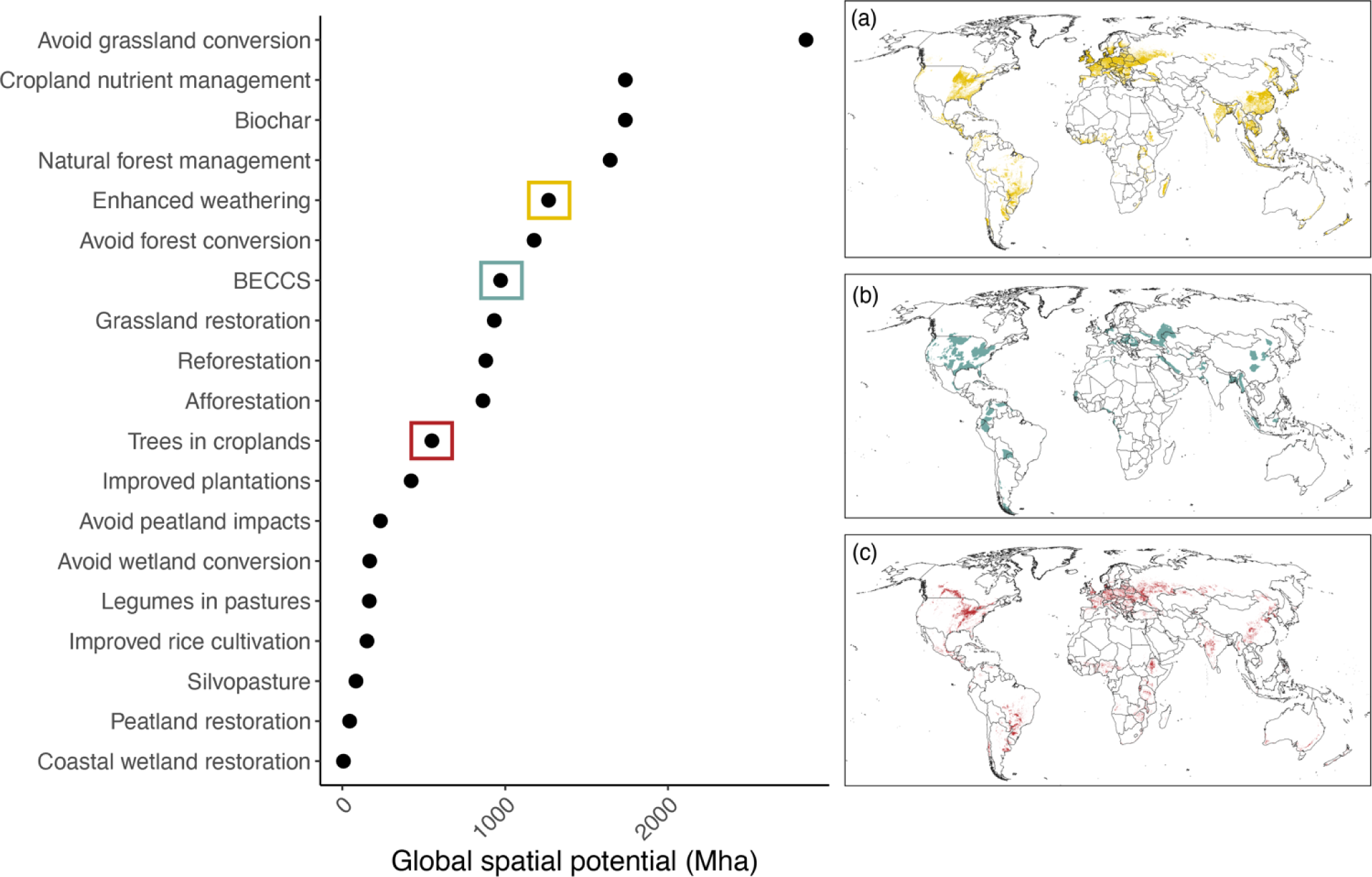
Spatial potential for 19 land-based climate mitigation strategies. The global land area available for implementing each mitigation strategy, in millions of hectares. Example distributions are shown by the inset maps for (a) enhanced chemical weathering, depicting croplands, pastures, and plantation forests within suitable environmental conditions, (b) bioenergy with carbon capture and storage (BECCS), depicting areas suitable for one of five common bioenergy crops that overlap with a saline sedimentary basin for CO_2_ storage, and (c) integrating trees in croplands, depicting croplands that have a high predicted potential for added tree cover.

### Global overlaps

After overlaying the spatial potential for each LBMS, we estimate that mitigation strategies could be applied to a maximum of 8.9 billion ha of Earth, equal to ∼60% of the global land area (Figure 2). This includes 3.3 billion ha of overlapping LBMS (i.e., more than one mitigation strategy is possible) and 5.6 billion ha of non-overlapping LBMS (i.e., only one of the analyzed LBMS could be applied).

**Figure 2.**
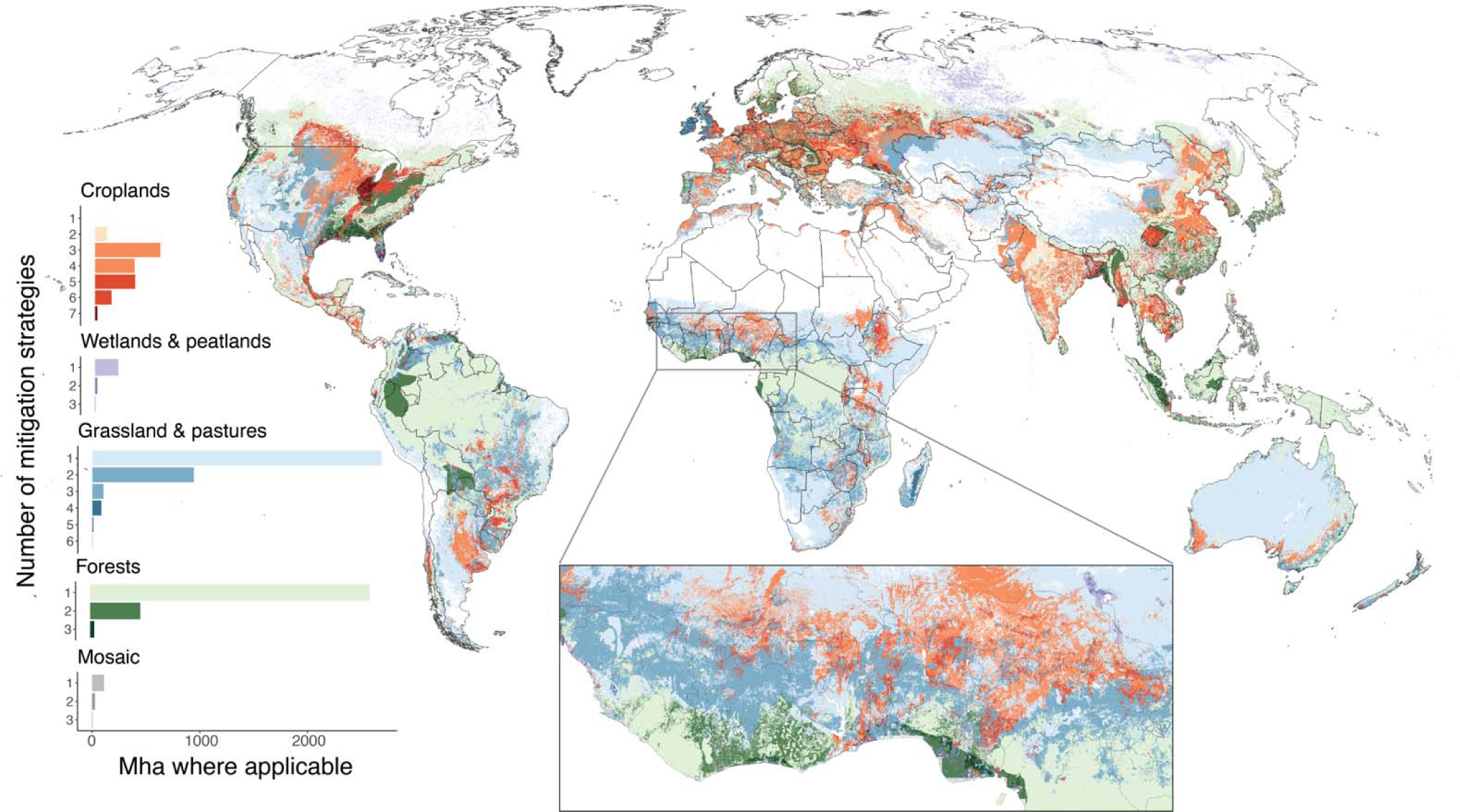
Global distribution of the land available for climate mitigation strategies. Spatial heterogeneity in the number of land-based mitigation strategies (LBMS) that are possible in any one area, delineated by the current land cover type. Land cover types include croplands, wetlands and peatlands, grasslands (including shrublands and savannas) and planted pastures, forests (including natural, managed, and plantation forests), and mosaic vegetation. Darker colors indicate a higher number of strategies overlap in that region. Legend bars represent the total area (in millions of hectares) where that number of LBMS could be applied in each land cover type, highlighting the large area of overlapping LBMS in croplands and pastures, and the non-overlapping area in forests, grasslands, wetlands, and peatlands. The inset map highlights the spatial variation and fine resolution (∼1km) of the land potential for climate mitigation in West Africa.

Given the non-random and differing distribution of individual LBMS, overlaps are also non-randomly distributed (Figure 2). For example, due to the large number of LBMS that can be implemented in agricultural settings (Table 1), croplands and pastures were most commonly identified as areas with multiple options for which LBMS to deploy. This includes for example the Great Lakes region of North America, croplands in China, and pastures in Madagascar, where up to six or seven LBMS overlap (Figure 2). This high degree of overlap includes multiple complementary management strategies that can be added to the same landscape. For example, up to 21% of croplands could integrate tree cover, receive enhanced weathering, and biochar (Figure S20). Up to 37% of pastures could be converted to silvopasture with enhanced weathering, optimal grazing, and legumes as cover crops (Figure S21). These croplands and pastures could also be converted to BECCS, afforestation, or restored to grassland.

Locations with three or more mitigation strategies are less common in forests, grasslands, wetlands, and peatlands (Figure 2 barplot). In forests, the maximum number of LBMS occurs in plantations suitable for enhanced weathering and bioenergy with carbon capture and storage (BECCS). This occurs for example in the southeastern United States, coastal Nigeria, and Europe (Figure 2). In grasslands excluding pastures, the areas with three or more strategies most often involve overlap between avoiding grassland conversion, BECCS, and afforestation (e.g., western United States, parts of Brazil, Africa south of the Congo Basin).

Excluding areas of overlap, we estimate 5.6 billion ha where only one of the analyzed LBMS could be applied. This area is overwhelmingly dominated by an opportunity to apply natural forest management (Figure 3 blue regions) or protect the carbon already stored in extant grasslands and tropical forests (i.e., avoiding carbon loss from historical rates of habitat conversion, Figure 3 pink regions). Ten additional LBMS have portions of land that occur in isolation of the other strategies, including a large spatial potential for reforestation in the tropics and subtropics and protecting wetlands and peatlands at northern latitudes.

**Figure 3.**
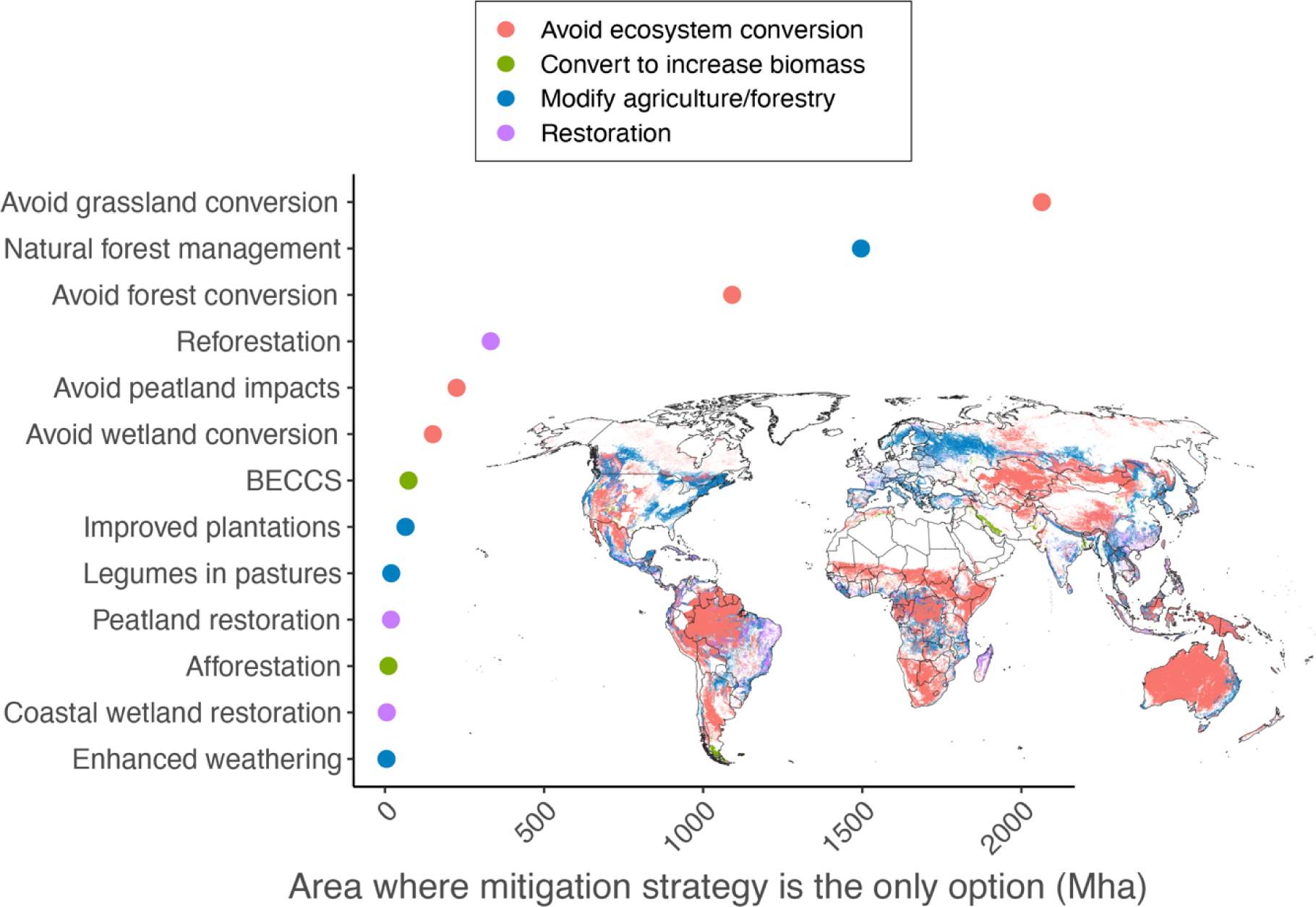
Mitigation strategies that occur in isolation of other LBMS. The global area (Mha) and distribution (inset map) of the 13 land-based mitigation strategies that occur within the area where only one of the studied strategies is possible (i.e., within the non-overlapping area of Figure 2). Colors denote which approach to climate mitigation applies (Table 1).

### Intersections between land-based mitigation strategies

To assess how scaling up the deployment of one LBMS might affect the area available to others, we estimated the absolute area and proportional overlap between each pair of mitigation strategies (Figure 4). We also considered where overlapping pairs of LBMS are mutually compatible, meaning they can both be deployed on the same land, or conflicting, meaning that the deployment of one strategy precludes the deployment of the other (SI). In the mutually compatible scenario, stakeholders may choose to apply one or both LBMS, while in the conflicting scenario, they must decide between strategies.

**Figure 4.**
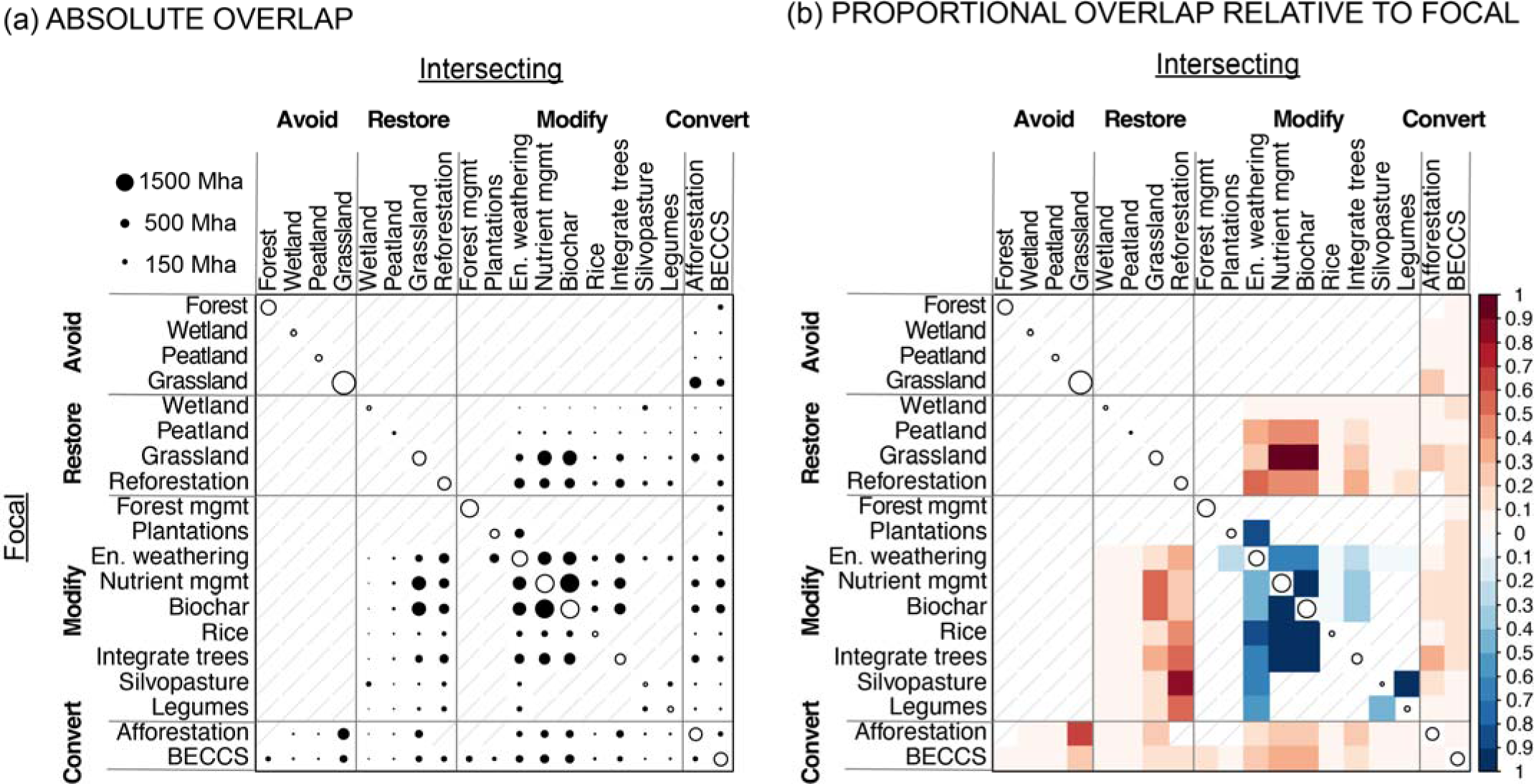
Pairwise overlap between land-based mitigation strategies. (a) The absolute area, scaled to Mha, where each pair of mitigation strategies overlaps across the globe. (b) Proportional overlap, measured as the area of overlap from panel (a) divided by the area suitable for the focal strategy listed in the rows (e.g., the uppermost blue square indicates that enhanced weathering overlaps with 84% of the area suitable for improved plantations). Because panel (b) presents intersections relative to the focal strategy, the plot is asymmetric across the diagonal. Blue intersections indicate LBMS that can both be deployed in the same area, whereas red intersections indicate LBMS that are incompatible with one another (SI). In both plots, the circles along the diagonal are scaled to the total area suitable for the focal strategy (Fig. 1) and grey lines indicate strategies that, by definition, do not overlap across space (e.g., reforestation cannot be applied to areas that are already forested). Strategies are grouped based on their approach to mitigation (Table 1) - avoiding ecosystem conversion, restoring ecosystems, modifying agriculture and forestry management, and converting land to bioenergy or afforestation.

The absolute and proportional overlaps vary depending on whether the mitigation strategy involves avoiding, managing, converting, or restoring land (Figure 4), the total extent available to each strategy in the pair, and their geographic co-variation. Conflicting LBMS account for the majority of intersections between strategies. The major conflict is between modifying the management of existing agricultural lands (‘Modify’ in Figure 4) and restoration (‘Restore’ in Figure 4). If biochar, for example, is to be deployed across its full extent, it would reduce opportunities for restoration by up to 48% for peatlands, 97% for grasslands, and 49% for forests. On the other hand, if reforestation is prioritized across its extent, it could significantly preclude opportunities for hybridizing agriculture via integrating tree cover into croplands (56%) or silvopasture (84%).

Converting land to bioenergy with CCS (BECCS) or afforestation intersects with all other LBMS, but typically only in the 10-30% range. However, these are the only strategies that intersect with avoiding ecosystem conversion, meaning that their deployment could convert land currently storing carbon in the natural state of the ecosystem. Although for forests, wetlands, and peatlands, the absolute and proportional intersections with BECCS and afforestation are small, for avoiding grassland conversion, the intersections are much larger (Figure 4A). In fact, if afforestation was fully realized, it would convert up to 21% of grassland area. And even though BECCS intersects with only 8% of the area suitable for avoiding grassland conversion (Figure 4B), this fraction represents 229 Mha of land (Figure 4A), an area twice the size of China.

Conversely, if avoided grassland conversion were prioritized over the intersecting LBMS, the area available for afforestation and BECCS would be reduced by 67% and 23% respectively (Figure 4B bottom rows).

Although conflicting pairs of strategies predominate, mutually compatible strategies were common among the various land management options (pairs within the ‘Modify’ category). Specifically, 13 pairs of mutually compatible strategies occur across LBMS that involve modifying cropland, pasture, and forest management (Figure 4B). For example, enhanced chemical weathering can be added to 84% of the area for improved plantations, 67% of areas where tree cover can be integrated into existing croplands, and up to 59% of areas suitable for sowing legumes in pastures. These proportional overlaps also represent large areas of Earth in absolute terms (Figure 4A), meaning that mitigation can result from multiple strategies for adapting agricultural and forest management over much of the Earth’s surface.

## Discussion

Reducing global climate change via a net zero emissions economy requires dramatic cuts to carbon emissions from the energy sector alongside increases in land-based climate mitigation^1,5,6^. Prior to this study, we lacked a synthesis of the land requirements and geographic constraints for the diversity of land-based approaches for reducing climate change. We addressed this gap by assembling 1) spatially explicit estimates of opportunities for implementing 19 LBMS across the globe and 2) key messages from those estimates, including cases in which scaling up one mitigation strategy might have consequences for the deployment of others. Importantly, we estimated almost nine billion hectares where land-based climate mitigation can be implemented across Earth’s terrestrial surface.

It is important to note that these data products are provided at a relatively high resolution (1 km) but the land suitability/mitigation potential is only constrained by biophysical or geographic variables. As such, our estimated spatial extents define the area within which additional factors, namely land tenure and socioeconomic attributes – many of which are highly uncertain^2,5^ - could determine the efficacy of mitigation. It is also important to note that we only consider mitigation strategies that rely on plant and soil processes to remove or offset emissions. Other demands on land for reducing climate change will come from technological solutions, such as the deployment of solar and wind energy, but these are beyond the scope of what we analyze here. By focusing on mapping the choices society faces for scaling up land-based climate mitigation, our analysis clarifies which strategies can be deployed across the globe and when overlapping strategies complement or conflict with one another.

We found that land-based climate mitigation – including individual strategies and their overlaps – is spatially-biased given Earth’s non-random distribution of land-use types. For example, several mutually compatible LBMS can be implemented in agricultural areas, including in particular, croplands and pastures in the eastern United States, throughout Europe, and in southeast Asia. By contrast, in natural forests, wetlands, and peatlands avoiding habitat conversion is often the only feasible mitigation strategy. While, in principle, one could deploy other LBMS in these habitat types, the loss of naturally stored carbon from land-use change would rarely result in net carbon removals^34,35^. These habitat conversions are thus not considered LBMS in this paper. Indeed, at a global scale we find that avoidance based LBMS (alongside natural forest management) account for the majority of the area where only one mitigation strategy can be deployed. The need for development in many of these regions may challenge the degree to which their conservation can contribute to mitigation, particularly outside of areas that are already be protected (e.g., Amazon rainforest, indigenous lands in Australia)^36,37^. Even so, habitat conservation has multiple co-benefits^16,37,38^, and thus protecting existing carbon pools in the large area suitable for avoiding habitat conversion should be a top priority for reducing climate change.

Global opportunities for restoration are also key to reducing climate change, and here we provide spatial estimates for where restoration could be implemented to restore forests, grasslands, peatlands, and coastal wetlands (Figure S14-17). Restoration has clear climate and biodiversity benefits^1,16^, but the scaling up of alternative mitigation strategies could displace opportunities for restoration. In particular, incentivizing novel cropland management strategies (e.g., enhanced weathering, biochar), and BECCS to a lesser degree, have the potential to reduce the land that could be restored (Figure 4B), though the economics of food production in these regions will likely be the deciding factor. In addition, we were not able to map mitigation potential from better management of rangelands (SI), and thus some of the area we map as suitable for restoration likely further conflicts with rangeland management. Most climate mitigation scenarios expect some reduction in the extent of land used for agriculture in favor of restoration^6^, and here we show where that could be implemented, including reforestation throughout China and Brazil, peatland restoration in northern Europe, and coastal wetland restoration in Central America.

Together, our results highlight important choices that governments and local communities must make in order to scale up land-based climate mitigation. We identify some opportunities for LBMS to work in concert, especially on agricultural land, but we find that LBMS are often in conflict with one another. A key take-home from our analysis is that increasing the implementation of a particular LBMS, such as policy initiatives that encourage carbon capture and storage^6,39,40^, should carefully consider the opportunity cost of reducing the land available to alternative approaches (e.g., better land stewardship). These opportunity costs may be unavoidable in many locations and will certainly have broad implications for not only climate change, but also for global biodiversity and human livelihoods.

To our knowledge, the collection of maps we have synthesized are derived from the best available data on the geography of land-based climate mitigation, and therefore serve as an important resource for coordinating land allocation to LBMS. Nonetheless, several limitations are worth noting and provide room for further exploration. For example, greater uncertainties are associated with emerging LBMS (enhanced chemical weathering, BECCS) and these are therefore mapped at a coarser spatial resolution^41,42^. The area available to restoration and to several of the cropland management strategies is likely to be overestimated given a lack of data on current land-use practices^20,22,43^. More generally, remote sensing products tend to underestimate the extent of certain land cover types used to map LBMS, such as wetlands, pastures, and other types of grazing lands^44,45^. This would lead the current study to underestimate the potential of strategies applied to these lands. At fine spatial scales, the maps may be sensitive to the limitations of underlying datasets and the rules we used to define suitability (SI). However, the sources we used are based on well-validated remotely sensed products, empirical data, and theoretical models ^27,45,46^, and we expect the spatial patterns and broader take-homes to be robust to minor changes in the input layers.

Finally, we focused on mapping the area where LBMS could *possibly* be implemented, and conflicts/complementarities could *possibly* arise. Future work is needed to estimate the probability of different outcomes across LBMS, which likely depends more on political and socioeconomic factors than the biophysical environment^6,8^. Nonetheless, our analysis provides a key step to estimating how much land is actually needed for LBMS to mitigate climate change, with the hope that these data products can guide future studies on optimizing pathways for reducing climate change.

## Methods

We compiled a list of 24 land-based climate mitigation strategies (LBMS) from the literature, defined as approaches that harness the processes that occur in vegetation, soils, and ecosystems to reduce carbon emissions and/or increase carbon sequestration (Table 1, SI). This list includes strategies previously defined as natural, nature-based, novel, and technological climate mitigation strategies/solutions, all of which modify land processes to alter carbon cycling and storage and could be deployed at large spatial extents to help reach net zero emissions.

Each land-based mitigation strategy, described in Table 1, has a continuous land footprint and is associated with a specific land cover type and/or set of geographical and biophysical requirements for where it can be implemented. This includes 18 of the 20 ‘natural climate solutions’ as defined and described by Griscom et al. (2017)^1^; we excluded fire management due to uncertainties in the spatial extent and carbon benefits of this as a management practice, and we excluded reduced harvest of woodfuel for cooking and heating because the climate benefits are estimated as a function of the number of people who abandon this practice, not the area where it applies. We also considered the land potential of seven other climate mitigation strategies not discussed by Griscom et al. (2017), including regenerative annual cropping^47^, silvopasture^48^, grassland restoration^20^, afforestation^49^, enhanced chemical weathering^50^, and bioenergy crop production paired with carbon capture and storage (BECCS)^18^.

Five of the mitigation strategies have specific geographic requirements but could not be mapped due to insufficient spatial data (SI). This included regenerative annual cropping, conservation agriculture, optimal grazing, improved animal feed, and improved animal management. We chose to retain these in Table 1 because previous studies have reported the maximum area available for each strategy^1^, and most of these are complementary to those which we were able to map (e.g., optimal grazing and improved animal feed/management complements other grassland conservation and management strategies, SI). For all spatial analyses, we did not include these five LBMS.

### Map derivation

For the remaining 19 LBMS, we identified and mapped their global distributions based primarily on geographic features and the biophysical environment. We assume that mitigation strategies will be deployed in the near-term, and thus maps are based on earth’s current distribution of land and climate (i.e., we do not consider future projections of land-use or climate change and their influence on where LBMS could be deployed, nor do we consider how the demand for LBMS might decrease given societal changes, such as an increase in vegetarian diets or renewable energy efficiency). All spatial data were compiled or derived from existing datasets, which include remotely sensed distributions of land cover types (e.g., Copernicus land-cover products)^51^, historical land cover changes^21,52^, and predictions of suitability based on contemporary geographic and climatic conditions^41,42^. All maps are binary, resampled to the same ∼1 km resolution, and harmonized to the International Union for Conservation of Nature (IUCN) habitat classification scheme^45^. Data sources, map derivation, and validation are described briefly below and in detail in Table S1 and SI.

Strategies that reduce business-as-usual carbon emissions by avoiding ecosystem conversion (Table 1) were primarily mapped using current distributions of habitat types as defined by Jung et al. (2020)^45^. We also used this dataset to identify croplands and pastures suitable for grassland restoration, biochar, cropland nutrient management, sowing legumes in pastures, and improved rice cultivation^53^. For natural forest management and improved plantations, we used global forest management data to classify different intensities of forest management^54^. For integrating trees in croplands and silvopasture, we combined Jung et al. (2020) with a predicted layer of global tree potential^55^, identifying croplands and pastures with high potential for added forest cover. We also used global tree potential^55^ and a global map of biome types^46^ to identify areas suitable for reforestation (restoring tree cover in forested biomes) and afforestation (adding tree cover to historically open biomes). Maps of peatland and coastal wetland restoration were derived from global products on remotely sensed land cover change^21,22,52^.

Maps of enhanced weathering and BECCS are based on theoretical environmental potentials and some infrastructure constraints, but we acknowledge that additional technical limitations will play a significant role in where each of these strategies will ultimately be placed^56^. We mapped enhanced weathering in areas with a theoretical potential for carbon capture given temperature and aridity^41^, and restricted these areas to managed landscapes where the habitat structure is suitable for mechanically spreading the minerals (croplands, pastures, planted forests). To map BECCS, we integrated data on predicted bioenergy crop yields^42^ and maps of sedimentary basins identified as high priority for carbon capture and storage (CCS)^57,58^. It is possible that bioenergy crop production is implemented without CCS^6,59^ or that new transmission pipelines could transport CO_2_ over long distances to reach an injection site^58^. However, without CCS, the mitigation potential of bioenergy drops substantially^59^, and we currently lack the infrastructure for cost-effective, long-distance CO_2_ transport^58^. We therefore restrict bioenergy potential to areas that overlap with sedimentary basins suitable for CCS, assuming that within these regions, CO_2_ from bioenergy crops could be sustainably transported to an injection site^58^.

At finer spatial scales, the maps are inherently sensitive to the decisions we used to derive them (e.g., a lower tree cover threshold for reforestation increases the area for this LBMS). However, the rules we used are dictated by well-resolved remotely sensed data products and well-validated models. We therefore expect large-scale patterns to hold regardless of details of individual datasets, and we provide all source data and code such that users can explore sensitivity to the decisions used to discretize maps (SI).

### Spatial data analysis

We estimated the total spatial extent (in millions of hectares, Mha) and binary distribution for where each strategy could be implemented across the globe. Using these distributions, we estimated the spatial overlap among all, and between each pair of LBMS. For the first overlap analysis, we stacked the 19 maps of mitigation strategies to estimate the total number of strategies suited to the geography of each 1km grid cell across the globe. The primary geographic constraint on each strategy was the land cover type where it can be applied (e.g., biochar only applies to croplands, improved plantation rotations only applies to planted forests under timber production), and thus the maximum number of deployable LBMS – and the strategy options – differ depending on the current land cover type. We thus quantified the possible number of LBMS conditioned on whether the strategy falls in an area that is currently defined as a cropland, forest (including natural, managed, and plantation forests), wetland, peatland, pasture, or grassland (including shrublands and savannas). Areas outside of these land cover types are part of the built environment, deserts, or small fractions of mosaic vegetation^45^.

To quantify the intersections between LBMS, we identified the absolute and proportional overlap between each pair of mitigation strategies. The absolute area is the total land area that is geographically suitable for both members of the pair. We also estimated proportional overlap between strategies as follows. For a given LBMS, we divided the absolute area of its overlap with another strategy by the total area over which the focal strategy could be applied. Doing so allowed us to assess how much an intersecting solution overlaps the total area suitable for a focal solution.

To assess potential trade-offs, we also classified each pair of overlapping strategies as mutually compatible (both can be applied to the same landscape, with roughly additive carbon benefits) or conflicting (mitigation strategies cannot be applied to the same land cover type due to incompatible infrastructure or management needs). Given that each map we provide bounds the area where a mitigation strategy could feasibly be deployed, we assume that overlaps bound the area where conflicts and compatibilities could arise. Many pairs of mitigation strategies do not overlap by definition, given that 1) we consider current land cover types to be mutually exclusive across space (i.e., habitats are assigned as either forest, grassland, wetland or peatland), and 2) we assume these land cover types are static in the short-term (e.g., we do not allow for cropland LBMS to be deployed outside of existing agricultural areas). Our process for defining pairs as non-overlapping, compatible, or conflicting is described in SI.

### Source data

We have provided source data for each figure, access notes for all input sources (Table S1), and output maps of each mitigation strategy, which are publicly available online: https://doi.org/10.6084/m9.figshare.24933312. Input layers and code will be made available upon publication and by request to the corresponding author.

## Acknowledgements

This work was supported by Princeton University’s Carbon Mitigation Initiative funded by bp. We thank Stephen Pacala, members of Princeton’s Net-Zero America project, the Wilcove lab, and the Levine lab for helpful feedback throughout project development. We also thank the authors of all source data used to derive the maps presented here, with special thanks to Matteo Bertagni, Jean-Francois Bastin, and Lorenzo Menichetti for personal communications regarding data.

