## Supplementary material for "Global spatial potential for implementing land-based climate mitigation": SI

The PDF includes:	Page

Table S1. Data sources for input layers	2

Supporting methods	4

*Mitigation strategies with insufficient spatial data*	4

*Map derivation by mitigation strategy*	5

*Carbon flux estimates*	10

*Pairing compatible and conflicting strategies*	12

Supporting references	14

Table S1. Data sources for input layers. Derived maps for each mitigation strategy are available online: https://doi.org/10.6084/m9.figshare.24933312.

| Layer | Reference | Access link | Access notes |
| --- | --- | --- | --- |
| Current land cover types (base layers) | Jung et al. (2020)^1^ | https://zenodo.org/records/4058819 | Downloaded ‘lvl1_frac_1km_ver004.zip’ on 9/21/2023. Defines current distribution of land cover types (coded to be mutually exclusive): forests, grasslands, wetlands, croplands, pastures, plantations, mosaic vegetation. |
| Biomes and climate zones | Dinerstein et al. (2017)^2^ | <http://ecoregions.appspot.com/> | Downloaded shapefile 6/26/2023. Defines forested and non-forested biomes, used to differentiate reforestation, afforestation, and grassland restoration. Also used to define tropical, subtropical, and temperate zones. |
| Forest management | Lesiv et al. (2022)^3^ | https://zenodo.org/records/5879022 | Downloaded ‘FML_v3-2_with-colorbar.tif’ on 9/22/2023. Delineates unmanaged, managed, and plantation forests. |
| Peatland extent and restoration potential | Leifeld & Menichetti (2018)^4^ | Shared via personal communication | Accessed 7/5/2022. Defines existing peatlands and areas for peatland restoration. |
| Mangrove extent and restoration potential | Bunting et al. (2022)^5^ | https://zenodo.org/records/6894273 | Downloaded ‘gmw_v3_f1996_t2020’ on 8/22/2023. Defines areas for mangrove restoration based on historical loss data, as one component of coastal wetland restoration. |
| Salt marsh extent and restoration potential | Campbell et al. (2022)^6^ | https://daac.ornl.gov/cgi-bin/dsviewer.pl?ds_id=2122 | Downloaded ‘sm_loss.tif’ on 8/22/2023. Defines areas for salt marsh restoration based on historical loss data, as one component of coastal wetland restoration. |
| Rice distribution | Monfreda et al. (2008)^7^ | http://www.earthstat.org/ | Downloaded ‘HarvestedAreaYield175Crops_Geotiff’ on 5/17/2023. Defines areas under rice cultivation. |
| Potential forest cover | Bastin et al. (2019)^8^ | https://www.research-collection.ethz.ch/handle/20.500.11850/350258 | Downloaded ‘Total_potential.tif’ on 9/25/2023. Defines areas suitable for tree cover (reforestation, afforestation, silvopasture, integrating trees in croplands). |
| Current forest cover | Hansen et al. (2013)^9^ | http://earthenginepartners.appspot.com/science-2013-global-forest | Downloaded ‘2000 Percent Tree Cover’ on 9/25/2023. Defines areas that already have >30% tree cover, used to exclude already forested regions from reforestation, afforestation, silovpasture, integrating trees in croplands. |
| Enhanced weathering distribution | Bertagni & Porporato (2022)^10^ | Shared via personal communication | Accessed 1/31/2023. Defines areas suitable for enhanced chemical weathering. |
| Carbon capture and storage basins (USA) | NETL, NATCARB Atlas Saline Basin 10km Grid (open access) | https://edx.netl.doe.gov/dataset/natcarb-atlas-saline-basin-10km-grid | Downloaded ‘natcarb_saline_poly_shapefile.zip’ on 9/25/2023. Defines U.S. saline sediment basins for carbon capture and storage (BECCS). |
| Carbon capture and storage basins (global) | U.S. Geological Survey World Petroleum Assessment 2000 (open access) | https://certmapper.cr.usgs.gov/data/apps/we-data/ | Downloaded ‘tps_geog.e00’ shapefile on 2/9/2023. Defines global (outside of U.S.) saline sediment basins for carbon capture and storage (BECCS). |
| Potential bioenergy yield | Li et al. (2020)^11^ | https://zenodo.org/records/3274254 | Downloaded ‘Bioenergy_crop_yields.zip’ on 7/9/2019. Defines areas with a positive predicted bioenergy crop yield. |
| World countries | Esri Data and Maps, World Countries Generalized (open access) | https://hub.arcgis.com/maps/esri::world-countries-generalized | Downloaded 7/6/2022. Defines the study area of terrestrial Earth. |

**Supporting methods**

*Mitigation strategies with insufficient spatial data*

Of the 24 different LBMS in Table 1, we were unable to derive maps for regenerative annual cropping, conservation agriculture, optimal grazing, better animal feed, and better animal management.

Regenerative annual cropping and conservation agriculture both apply to most annual cropping systems^12^, but delineating annual from perennial cropping systems using remotely sensed data is challenging due to inconsistencies in crop spatial data, crop rotations, timing of harvest, and global cropland statistics^7^. Optimal grazing, improved animal feed, and improved animal management are specific to rangelands with livestock grazing^12,13^. However, we were not able to find a global, spatially-explicit dataset that delineated rangelands from low intensity or ungrazed grasslands, shrublands, and savannas. Grazing, as a mobile and periodic land management strategy, is difficult to detect via remote sensing^14,15^, and as a result, estimates of the global extent of rangelands vary substantially.

*Map derivation by mitigation strategy*

Derived maps for each mitigation strategy are available online: https://doi.org/10.6084/m9.figshare.24933312.

Base layers

We first defined Earth’s current distribution of land cover types using the International Union for Conservation of Nature (IUCN) habitat classification scheme^1^. This dataset defines terrestrial habitat types by integrating several land-cover datasets, including the Copernicus land-cover product^16^, the global Köppen-geiger climate classification system^17^, maps of terrestrial ecoregions^2^, among others. Data layers are 1km resolution estimates of the percent cover of different land cover classes. To define mutually exclusive land cover types, we assigned pixels with >50% of a single land cover type as one of cropland, grassland (including shrubland and savanna), pasture, unmanaged forest, managed forest, plantation forest, wetland, peatland, or mosaic vegetation (>50% of some combination of the former land cover types).

These classes helped define suitability for mitigation strategies as well the degree of land conversion that would result from implementing a strategy. All subsequent mapping was harmonized to the distribution, resolution, and native coordinate system of land cover types in Jung et al. (2020)^1^: ~1km World Geodetic System 1984. Any dataset that was not provided at this resolution or coordinate system was resampled and projected to match Jung et al. (2020) using bilinear interpolation. All maps were then discretized (1/0, suitable/unsuitable) using rules described below. To estimate the area available to each solution and overlaps, we extracted the identity of strategies suitable in each pixel and the pixel cell size. We then grouped by strategy and summed across pixels. Map derivations and analyses were done using the ‘terra’ package in R^18^. The final set of maps were masked to terrestrial Earth using ESRI’s World Countries shapefile (Table S1).

Avoided conversion of forests, grasslands, wetlands

Avoiding the conversion of forests, grasslands (including shrublands and savannas), and wetlands (including mangroves) applies to extant ecosystems, which were mapped using the classifications from Jung et al. (2020) described above. For avoiding forest conversion, we focused on forested pixels without signs of management^3^. Following Griscom et al. (2017)^12^, we restricted avoiding forest and grassland conversion to ecosystem types in temperate, subtropical, and tropical climate zones. Further, for avoiding grassland conversion, we only retained pixels in non-forested biomes^2^, assuming these represent historically intact natural grasslands, whereas grasslands in forested biomes likely reflect the loss of forest due to human land-use^9^.

Avoided peatland impacts, peatland restoration

To map avoided peatland impacts, we used a recent study by Leifeld & Manichetti (2018)^4^, which maps peatlands at a 1km resolution. Each pixel has an estimated ratio of intact to degraded peatland, ranging from 0 (fully intact) to 1 (fully degraded). We defined avoided peatland impacts as wetland pixels^1^ with a value ≤ 0.5 (mostly intact) and peatland restoration as pixels > 0.5 (mostly degraded).

Coastal wetland restoration

Coastal wetland restoration includes restoring degraded salt marsh and mangrove ecosystems. To map areas for restoration, we used two recent studies that analyzed global aerial imagery to estimate the extent of salt marsh change from 2000-2019^6^ and mangrove change from 1996 to 2020^5^. Because the study periods are only about two decades, both studies underestimate the total extent of wetland loss relative to earlier time stamps (e.g., Macreadie et al. 2021^19^ suggest a restoration potential of 0.2–3.2 Mha for tidal marshes and 9–13 Mha for mangroves). However, these represent the best available data at a global extent and a fine resolution of ~30 meters. For both datasets, we estimated the percent of habitat loss at ~1 km resolution over the study period. We then used a threshold of >1% mangrove loss and >25% salt marsh loss to classify areas as suitable for restoration. These thresholds were selected to result in a total area of restoration potential that fell towards the conservative end of the range of total area estimates suggested by previous studies: ~9 Mha of mangrove and ~0.3 Mha of salt marsh restoration^12,19,20^. There was a small degree of overlap between coastal wetland restoration and peatland restoration, in which we assigned pixels as peatland restoration given the higher carbon storage potential in peat soil (Table 1 in the main text).

Natural forest management, improved plantations

To map natural forest management and improved plantations, we used Lesiv et al. 2022^3^ data on global forest management to delineate forests into classes based on different degrees of management. Management classes were identified at a 100m resolution, which we aggregated to the percent of each class within a 1km pixel and resampled to the resolution of Jung et al. (2020)^1^. We considered natural forest management to apply to naturally regenerating forests with signs of management and planted forests with a long rotation time (>15 years). We selected pixels with >50% cover of naturally regenerating and/or planted forest and assigned this class to forested pixels from Jung et al. (2020). This identified 37% of forests with low-intensity management^3^.

For improved plantations, we considered this strategy to apply only to the subset of planted forests that are intensively managed for timber production, defined as planted forests with a short rotation time (<15 years)^3^. We followed the same process of aggregating the data to 1km, resampling using bilinear interpolation, and selecting pixels with >50% cover of plantations. As expected, most of the plantations defined by Lesiv et al. (2022) were already classified as such by Jung et al. (2020), but the latter defined ~2.5 Mha of plantations as other types of forest. We harmonized the data sources to a single layer of plantations, including Jung et al. (2020) mapped plantations and Lesiv et al. (2022) defined plantations that fell within classified forests.

Biochar, cropland nutrient management, improved rice

Adding biochar and improving cropland nutrient management could theoretically apply to all croplands where these techniques are not already used. To map croplands where these solutions could apply, we selected pixels with >50% cover of arable land^1^. Recent critiques have argued that assuming these mitigation strategies apply to all croplands is an overestimate of potential^21,22^. While the magnitude of carbon sequestration likely varies spatially, to our knowledge, there is no known geographic limitation to improving the carbon storage of agricultural soils. We therefore considered biochar and cropland nutrient management to be possible on any extant cropland.

To map improved rice cultivation, which can theoretically apply to anywhere that rice is presently cultivated^12^, we used crop maps generated by Monfreda et al. (2008)^7^. These data products estimate the hectares harvested for different crops at a 10km resolution across the globe. For each grid cell, we estimated the proportion of hectares harvested of rice relative to the hectares harvested of non-rice crops. We resampled the data to a 1km resolution using bilinear interpolation and assigned cropland pixels as rice if >25% of the harvested area could be attributed to rice cultivation. We selected this threshold to assign enough area to match global expectations of ~165 million hectares of rice cultivation^12,23^. We verified that assigning this distribution of rice matched expectations for highly productive regions of rice production (e.g., China, India, Indonesia).

Trees in croplands, silvopasture

To identify areas where tree cover could be integrated into croplands (excluding ricelands) and pastures (silvopasture), we used Bastin et al. (2019)^8^ global layer of total tree potential. This layer is an unconstrained estimate of Earth’s tree carrying capacity predicted at a 1km resolution based on empirical observations of trees in protected areas. Following well-validated rules for defining forest using remotely sensed data products^9^, we identified pixels as suitable for increasing tree cover if they had >30% predicted tree cover^8^ and <30% current tree cover^9^, resulting in a layer of croplands and pastures with low tree cover that are suitable for increases.

Enhanced weathering

To map potential areas for enhanced chemical weathering, we used Bertagni & Popporato (2022)^10^ rules for constraining the theoretical potential for enhanced weathering (without irrigation) to any biome with an average temperature > 0°C and a global aridity index > 3 (warmer and wetter biomes). We considered any area with a positive carbon capture efficiency rate^10^ to be suitable for enhanced weathering, but limited enhanced weathering to managed systems where the infrastructure might be available to spread the minerals (croplands, pastures, plantation forests).

Legumes in pastures

Sowing legumes in pastures could theoretically be applied in all planted pastures where this technique is not already used. To identify pastures as suitable for legumes, we selected pixels with >50% cover of pastureland as defined by Jung et al. (2020).

Reforestation, afforestation

Following a similar process for mapping trees in croplands and silvopasture, we defined suitable areas for reforestation and afforestation as non-forested pixels with high suitability for increased forest cover^1,8,9^. We identified reforestation potential as pixels within forested biomes and afforestation potential as pixels in non-forested biomes^2^, thus delineating areas where historical forest could be restored from areas where adding tree cover would convert the landscape to a novel state. We excluded reforestation and afforestation potential in the boreal zone because such efforts are generally thought to warm rather than cool the climate by altering albedo. We also removed any reforestation potential in areas already assigned as suitable for peatland or coastal wetland restoration.

We elected to derive our own reforestation and afforestation layers rather than use existing maps of reforestation potential to avoid any prescribed limits based on uncertain socioeconomic factors. For example, the map of reforestation potential by Griscom et al. (2017)^12^ avoids reforestation in certain croplands to protect food security; the World Resources Institute Global Restoration Initiative layer (https://www.wri.org/initiatives/global-restoration-initiative) avoids areas with high intensity human land-use; Bastin et al. (2019) received criticism for not separating reforestation potential (restoring historically forested areas) from afforestation potential (adding trees to systems that were not historically forested) (eLetter responses^8^). Because we are exploring the mitigation strategies that are possible, not the choices that are most probable or least controversial, we derived an unconstrained prediction of potential for reforestation/afforestation, as well as for other mitigation strategies that integrate trees into agricultural areas (e.g., trees in croplands, silvopasture).

Grassland restoration

We defined grassland restoration as possible in any cropland or pasture in a grassland biome^2^, assuming these areas were historically natural grasslands that were converted to agriculture (we assume that outside of open biomes, grassland habitat represents degraded forest which would be suitable for reforestation rather than grassland restoration). We exclude grassland restoration in areas already assigned as suitable for peatland or coastal wetland restoration. Other areas are likely suitable for grassland restoration (e.g., degraded or overgrazed grasslands, and grasslands dominated by non-native, invasive species) but are more difficult to differentiate from intact natural grasslands using only remotely sensed data products.

Bioenergy with carbon capture & storage (BECCS)

To identify areas that could be used for BECCS, we combined maps of predicted bioenergy crop yields^11^ and maps of sedimentary basins identified as high priority for CO_2_ storage^24,25^. For the bioenergy yield maps, Li et al. (2020)^11^ compiled globally distributed empirical data on yields of five common bioenergy crops: Miscanthus, switchgrass, willow, poplar, and eucalyptus. They predicted maximum yield across the five crop species at ~50 km resolution, which is notably coarser than the other data sources used in this analysis, due to limited empirical data on bioenergy crop yields for many regions of the Earth. But to our knowledge, the analysis provided by Li et al. (2020) uses the best available empirical data on bioenergy crop production and avoids sensitivities associated with socioeconomic assumptions embedded in other bioenergy predictive modeling frameworks^26^ (e.g., integrated assessment models). The dataset is therefore comparable with other maps used in the analysis and represents a coarse prediction of the areas that are suitable for bioenergy crop production. We resampled the data to a 1 km spatial resolution using bilinear interpolation and considered anywhere with a positive predicted yield as suitable for bioenergy crop production.

It is possible that bioenergy crop production is implemented without CO_2_ storage, in which case the range of possible locations where bioenergy could occur is larger than what we present here. However, as stated in the main text, without CCS, the mitigation potential of bioenergy drops substantially^27^, and we currently lack the infrastructure for long-distance CO_2_ transport^24^. We therefore restricted bioenergy to only locations that overlap with a suitable sediment basin for carbon storage. For the latter, we used the U.S. Geological Survey U.S. & 2000 World Assessment of Petroleum Resources maps of high priority saline sedimentary basins (Table S1). These maps provide the boundaries for sedimentary basins with high potential for carbon storage. We assumed that an injection site and facility could be located anywhere within the boundary of a basin, and thus a bioenergy cropland located within the basin boundaries would be within a distance where carbon could be sustainably transported to the injection site. We therefore identified areas as suitable for BECCS if a pixel had a positive predicted bioenergy yield and overlapped with a suitable carbon storage basin.

*Carbon flux estimates*

In Table 1, we include a rounded carbon flux estimate for each mitigation strategy. Following Griscom et al. (2017)^12^, these estimates coarsely capture variation in the intensity of carbon emissions avoided or additional carbon sequestered from the implementation of each strategy, per hectare per year. These estimates are only intended to provide a sense for how strategies compare in the order of magnitude of mitigation potential, which undoubtedly varies across space, time, and with various management and technical constraints. Further, these estimates correspond to different types of carbon gains (emissions avoided vs. additional carbon sequestered) and so may not be directly comparable across strategies.

For the 18 mitigation strategies described by Griscom et al. (2017), we rounded the intensity of carbon flux values reported in their Appendix Table S1. When applicable, we extracted the estimates that apply at the global scale rather than estimates specific to each climate zone. For coastal wetland restoration, Griscom et al. (2017) reports a value for avoidable flux and additional carbon sequestered. We extracted the value for avoidable flux (to be consistent with the value for peatland restoration). Most values were already reported in carbon or carbon equivalents per hectare, but for several mitigation strategies, we converted the original units as reported below. For the mitigation strategies not discussed in Griscom et al. (2017), we followed a similar process for estimating carbon per hectare per year.

Cropland nutrient management: Units for carbon flux were reported as carbon equivalents (Ce) per Mg of Nitrogen fertilizer^12^. To estimate the Ce per hectare, we assumed a constant rate of nutrient application of 63.2 kg N per hectare (median estimate across the world, reported in Table S4 of Menegat et al. 2022^28^). We converted this to Mg of N and multiplied this by the estimated 4.33 Mg Ce / Mg N^12^. This resulted in 0.27 Mg Ce/ha/year avoided emissions from better cropland nutrient management, rounded to <1 Mg Ce sequestered.

Improved animal feed/management: Griscom et al. (2017) reports 0.13 Mg Ce sequestered per head of cattle, given 1400 M head of cattle globally. We assumed these cattle are distributed across 529 Mha of rangeland habitat suitable for improved grazing^12^, resulting in an average estimate of 0.11 MgCe/ha, rounded to <1 Mg Ce sequestered.

Bioenergy with carbon capture & storage: We estimated the carbon benefits of bioenergy with CCS as the carbon offset from reducing the consumption of fossil fuels and the carbon sequestered from increasing aboveground biomass^27^. We then reduced this estimate based on an assumed level of emissions that might result from fertilizer application to the crop system^28^ as well as emissions from the conversion of the original land cover type^29^. We arrived at a global range of 1-10 Mg C avoided and sequestered.

Grassland restoration: We took the median across continent-specific flux estimates reported in Figure 4 of Bai & Cotrufo 2022^30^. We rounded this estimate to <1 Mg C sequestered.

Conservation agriculture & regenerative annual cropping: Rounded median estimate across manure, cover crops, and reduced tillage, reported in Table 1 of Schlesinger et al. (2022)^31^.

Silvopasture: Rounded median estimate across observed silvopastoral systems in the temperate^32^ and tropical zones^33^.

Enhanced weathering: Rounded median estimate across global net carbon sequestration potential under a business-as-usual scenario reported in Beerling et al. (2022)^34^.

Afforestation: Rounded median estimate across the global carbon accumulation estimates for non-forested biomes, reported in Cook-Patton et al. 2020^35^.

*Pairing compatible and conflicting strategies*

There were many pairs of mitigation strategies that by definition, do not overlap across space. This is primarily a function of Earth’s current distribution of land cover types, which we considered to be mutually exclusive at the 1km scale. For example, habitat types are assigned as one of forest, wetland, grassland or peatland, so it is not possible to avoid the conversion of multiple habitat types in the same 1km grid cell. Similarly, agricultural areas are mutually exclusive (pasture, cropland, low-intensity managed forest, high-intensity planted forest), and thus strategies that apply to existing pastures do not apply to croplands and managed forests, and vice versa. We also do not allow for any of the agricultural management strategies to apply outside of areas that are already being used for agriculture. It is certainly possible for intact ecosystems to be displaced by agricultural expansion^36^, but given that our focus is on land conversions that benefit climate change, we exclude agricultural expansion outside of existing areas, with the exception of the potential of scaling up bioenergy with CCS.

Because we assume that avoiding habitat conversion applies to ecosystems that are already intact, we do not allow avoiding habitat conversion and restoration to overlap. Small scale restoration could potentially enhance the climate benefits of existing habitat types, but given that the maps are at a 1km resolution, we focus on large-scale restoration that we assume applies to degraded or converted habitat, given classifications in the underlying datasets (e.g., temporal data on global wetland loss^6^). Other than BECCS, we do not allow for any mitigation strategy to displace managed or plantation forests. This is to avoid losing carbon that is already stored in tree biomass, although we allow for the possibility of BECCS given that forestry byproducts are considered a potential source for bioenergy^37^. In which case, managed forests and plantations that overlap with CCS basins could be converted to BECCS.

For pairs of LBMS that do overlap, we assessed competition for space-use by defining each pair as mutually compatible (both can be applied to the same landscape) or conflicting (mitigation strategies cannot be applied to the same land cover type due to incompatible infrastructure or management needs). These definitions are also primarily based on the existing land cover type and the degree of land conversion that would result from applying a mitigation strategy. For example, the only pairs of mitigation strategies we consider to be compatible include those which modify the management of the existing land cover type (e.g., improving rotations in planted forests, soil amendments in croplands) and thus do not require a change in land cover type. One exception is integrating tree cover in areas where rice cultivation could be improved; we exclude this overlap because the latter refers to flooded rice systems, and it is unlikely tree cover can be added in areas that are periodically flooded and harvested.

Excluding the overlap among agricultural management strategies, all other pairs of overlapping LBMS are conflicting. We assume restoration includes restoring long-term above- and below-ground carbon pools, which is incompatible with continuing to sow and harvest biomass in croplands, managed forests/plantations, and pastures. At finer spatial resolutions, some amount of restoration could be compatible with agricultural management, but at the 1km scale, we consider restoration to be in conflict with any strategy that would require maintaining the land for large-scale agriculture or forestry. We also consider afforestation and BECCS to conflict with all other strategies, given they both convert land to a new state.
